# Structural Visualization of HECT-type E3 ligase Ufd4 Accepting and Transferring Ubiquitin to Form K29/K48-branched Polyubiquitination

**DOI:** 10.1101/2023.05.23.542033

**Authors:** Xiangwei Wu, Huasong Ai, Junxiong Mao, Hongyi Cai, Lu-Jun Liang, Zebin Tong, Zhiheng Deng, Qingyun Zheng, Lei Liu, Man Pan

## Abstract

The K29/K48-linked ubiquitination generated by the cooperative catalysis of E3 ligase Ufd4 and Ubr1 is an enhanced protein degradation signal, in which Ufd4 is responsible for introducing K29-linked ubiquitination to K48-linked ubiquitin chains to augment polyubiquitination. How HECT-E3 ligase Ufd4 mediates the ubiquitination event remains unclear. Here, we biochemically determine that Ufd4 preferentially catalyses K29-linked ubiquitination on K48-linked ubiquitin chains to generate K29/K48-branched ubiquitin chains and capture structural snapshots of Ub transfer cascades for Ufd4-mediated ubiquitination. The N-terminal ARM region and HECT domain C-lobe of Ufd4 are identified and characterized as key structural elements that together recruit K48-linked diUb and orient Lys29 of its proximal Ub to the active cysteine of Ufd4 for K29-linked branched ubiquitination. These structures not only provide mechanistic insights into the architecture of the Ufd4 complex but also provide structural visualization of branched ubiquitin chain formation by a HECT-type E3 ligase.

## Introduction

Poly-ubiquitination, formed by attaching a glycine at the C-terminus of one ubiquitin (Ub) to an amino group (M1, K6, K11, K27, K29, K33, K48, K63) on the surface of another Ub, governs almost all cellular processes in eukaryotes^1^. The functional diversity of poly-ubiquitination arises from the variable conformations of Ub chains, which can encode different functional signals to regulate the stability, localization and interactions of proteins, ultimately determining the fate of their substrates, known as Ub code^2^. Among these Ub codes, K48-linked Ub chains represent the best studied, acting as degradation signals that target substrate proteins to the proteasome^3,4^. Importantly, there is increasing evidence indicating that K48-linked Ub chains can be simultaneously ubiquitinated on other Lys (K) sites, leading to the formation of branched Ub chains^5,6^. The assembly of branched Ub chain could edit the original conformation of K48-linked Ub chain, thereby altering the functional signal. For example, the E3 ligase anaphase-promoting complex (APC/C) and E2 enzyme UBE2S assemble K11 linkages on K48-linked Ub chains K11/K48-branched Ub chains. These branched chains greatly enhance the proteasomal recognition of kinase Nek2A, thereby promoting the degradation of cell cycle regulators in the early stages of mitosis^7^. Induced by IL-1β signalling, E3 ligases HUWE1 and TRAF6 collaborate to assemble K63/K48-branched Ub chains, ultimately enhancing the NF-κB signaling^8^. Given the fundamental role of branched Ub chains in biological regulation, understanding how branched Ub chains are precisely assembled is one of the central concerns and challenges in the field.

The K29/K48-heterotypic Ub chains, first discovered in the N-terminal degradation pathway, were shown to accelerate the degradation of N-end substrates^9^. These heterotypic Ub chains are catalysed by two different E3 ligases, Ubr1 and Ufd4, of which the E3 ligase Ufd4 is responsible for augmenting K48-linked Ub chains in a K29-linked-specific manner^9^. Notably, a recent study has observed that E3 ligase TRIP12, the human homologue of Ufd4, can promote small-molecule-induced degradation through K29/K48-branched Ub chains^10^. However, the mechanism through which Ufd4 introduces K29-linked Ub chains on K48-linked Ub chains to increase the processivity of polyubiquitination remains unknown.

HECT-type E3 ligases are unique in their catalytic mechanism. Recent structural studies of the E3-E2-ubiquitin complex^11,12^, Ub-loaded E3 complex^13–15^ or E3-Ub-substrate complex^16–18^have illustrated the conformational plasticity at different stages of the reaction cycle. However, the mechanism by which HECT-type E3 enzymes assemble specific branched Ub chains remains unexplored. In this work, we biochemically demonstrate that Ufd4 preferentially catalyses K29-linked ubiquitination on K48-linked Ub chains, leading to the formation of K29/K48-branched Ub chains. By employing the biomimetic chemical probes to trap the enzymatic intermediates during the formation of branched Ub chains, we successfully determine the cryo-EM structures and reveal the mechanistic details of how the substrate Ub chains are recognized by Ufd4 to assemble the branched Ub chains. We also capture a structural snapshot of TRIP12 (the human homologue of Ufd4) transferring Ub onto K48-linked diUb, confirming the evolutionary conservation of these enzymes. In summary, our work showcases the inaugural structural visualization of HECT-E3 Ufd4 accepting and transferring Ub to form K29/K48-branched polyubiquitination.

## Results

### Ufd4 prefers to synthesize branched ubiquitin chains on K48-linked ubiquitin chains

Previous studies have shown that Ubr1/Ufd4 together strengthened the ubiquitination and degradation of the substrate (e.g., Scc1^9^ and the O^6me^G-DNA alkylguanine transferase Mgt1^19^) through K29-and K48-linked polyubiquitin chains, in which Ufd4 has been proposed to be a potential E4-like enzyme and act as a Ub chain elongation factor^20^. Accordingly, we began our study with the reconstitution of Ufd4-mediated polyubiquitination reactions on the substrates of monoUb, K29-linked diUb, or K48-linked diUb using the yeast Ub-activating enzyme (E1) Uba1, the Ub-conjugating enzyme (E2) Ubc4, the HECT-type E3 ligase Ufd4, and wild-type (WT) Ub. Enhanced polyubiquitination was observed on substrate K48-linked diUb compared to monoUb or K29-linked diUb (Fig. 1a and Supplementary Fig. 1a-g). The Ufd4 preference for K48-linked diUb was further confirmed by activity assays on eight different chain-type diUb substrates (M1, K6, K11, K27, K29, K33, K48 and K63 linkages) (Supplementary Fig. 1h). Moreover, Ufd4 ubiquitination assays on different lengths of K48-linked Ub chains (tri-, tetra-, and penta-) suggested that the polyubiquitination efficiency escalated with increasing chain length (Supplementary Fig. 1b).

**Figure 1.**
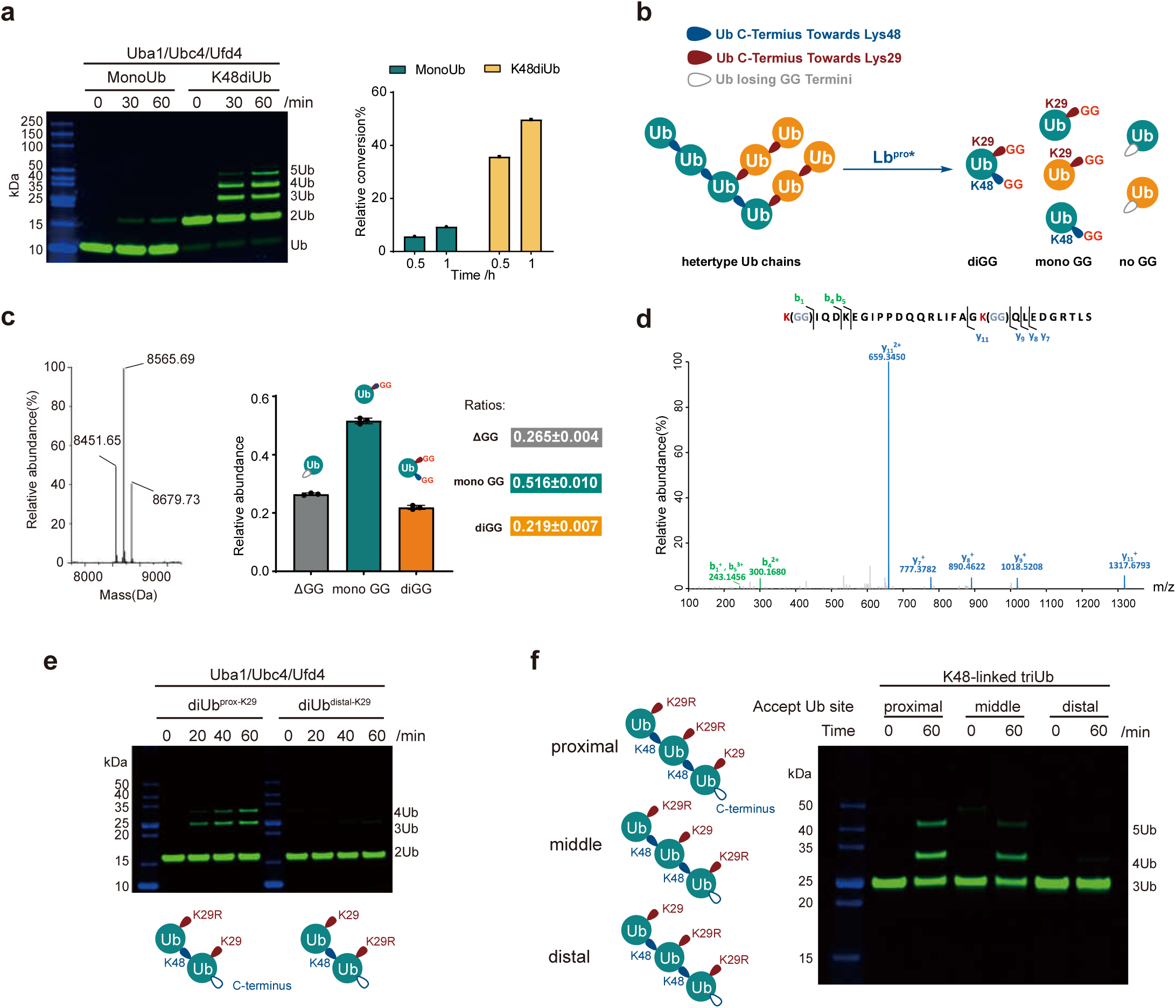
Ufd4 prefers to synthesize K29/K48-branched ubiquitin chains on K48-linked ubiquitin chains. **a**, Ufd4-dependent in vitro ubiquitination on fluorescent MonoUb and K48-linked diUb. Gel images and columns of relative conversions (%) are shown. Gel images are representative of independent biological replicates (n = 2). **b**, Schematic representation of the identification flow of Ub topology generated by Ufd4 on K48-linked Ub chains. The blue symbols represent the isopeptide bond between Lys48 and Ub C-terminal, the red symbols represent the isopeptide bond between Lys29 and Ub C-terminal, the grey symbols represent Ub C-terminals. **c**, Intact MS analysis of in vitro ubiquitination reactions treated with Lb^pro^*, and the spectra were deconvoluted. Deconvoluted spectra (left), quantification of the relative abundance of each ubiquitin species (right) are shown. Data represent the mean ± SD of three independent experiments. **d**, Identification of K29/K48-branched ubiquitin chains. The MS/MS spectrum corresponding to the signature peptide derived from K29/K48-branched linkages (aa 29–57, GlyGly modified at K29 and K48) was obtained. **e,** Ufd4-dependent in vitro ubiquitination on fluorescent K48-linked diUb with lysine to arginine mutation at the proximal or distal Lys29 sites. Gel images are representative of independent biological replicates (*n* = 3). **f,** The Ufd4’s ubiquitination activity on K48-linked triUb with a K29-only residue in the proximal, middle or distal Ub. Gel images are representative of independent biological replicates (*n* = 2). Source data are provided as a Source Data file.

Next, the topology of the Ub chains generated by Ufd4 on K48-linked Ub substrates was determined. First, our biochemical assay and mass spectrometric analysis revealed Ufd4 specifically mediates K29-linked ubiquitination on monoubiquitinated substrate (here, monoubiquitinated degron peptide, Supplementary Fig. 2a/b), in line with the previous results^21^. Meanwhile, the Ufd4-mediated polyubiquitination activity on K48-linked Ub chain substrates was significantly reduced when Ub-K29R mutant was used, supporting K29-linked ubiquitination on K48-linked Ub chain substrates (Supplementary Fig. 1c-f). Middle-down MS analysis (termed Ub-clipping^22^) of the polyubiquitination product (using K48-linked tetraUb as a representative substrate) revealed 21.9% of mono-Ub species modified by double-glycine remnants, representing the generation of branched Ub chains. Additionally, Ub fragments with double-glycine remnants on both K29 and K48 residues were detected in the MS/MS spectrum (Fig. 1b-d, Supplementary Note). To further elucidate the ubiquitination site preference of Ufd4, we performed ubiquitination assays using K48-linked diUb mutants with K29R mutation in either the proximal or distal Ub as substrates. The biochemical results demonstrated that mutants with proximal K29R mutations exhibited weak ubiquitination, whereas mutants with distal K29R mutations remained efficiently ubiquitinated (Fig. 1e). Furthermore, the enzyme kinetics of Ufd4 were determined individually for the aforementioned K48-linked diUb mutants and the results revealed that the ubiquitination efficiency, as represented by k_cat_/K_m_ (Supplementary Fig. 2c), is approximately 5.2-fold higher at proximal K29 site (0.11 μM^-1^·min^-1^) compared to distal K29 site (0.021 μM^-1^·min^-1^). We also prepared fluorescently labeled K48-linked triUb, with only one Ub (proximal, middle, or distal) retaining the K29 site, while the K29 sites in the other two Ubs were mutated to Arg (Supplementary Fig. 20) and then conducted Ufd4 ubiquitination experiments. Branched ubiquitination was observed on K48-linked triUb with the K29-only site in either the proximal or middle Ub. In contrast, under the same catalytic conditions, very weak ubiquitination activity was detected on K48-linked triUb with the K29-only site in the distal Ub (Fig. 1f). Taken together, these biochemical results suggest that Ufd4 preferentially catalyzes K29-linked Ub conjugates on preassembled K48-linked Ub chains to form K29/K48-branched Ub chains.

### Structural visualization of the K29/K48-branched ubiquitination by Ufd4

To investigate the molecular mechanism of how Ufd4 transfers its thioester-bound Ub to the proximal K29 of the K48-linked diUb substrate, we covalently linked the catalytic residue of Ufd4 (C1450), the C-terminus of Ub and the proximal K29 of K48-linked diUb to form a stable complex to mimic the corresponding transition state (Fig. 2). This complex was prepared in two steps: first, an engineered K29/K48-branched triUb probe (named triUb^probe^) was synthesized according to our previously reported procedure^23–28^, in which the proximal Ub of K48-linked diUb was chemically ligated with the donor Ub (Fig. 2a, b and Supplementary Fig. 3c,d); then, triUb^probe^ was cross-linked with Ufd4 in a catalytic residue (C1450)-dependent manner to form the designed stable complex mimic (Supplementary Fig. 3a). This complex was subjected to single-particle cryo-EM analysis. A total of 5,332 micrographs were collected and processed to obtain a cryo-EM map at a global resolution of 3.31 Å (Supplementary Tab. 1 and Supplementary Fig. 21). The AlphaFold-predicted Ufd4 structure (code: AF-P33202-F1) and the crystal Ub structure (PDB code: 1UBQ) were docked into the cryo-EM maps with meticulous adjustments and iterative refinements to obtain the model of the complex (Supplementary Fig. 4a, b).

**Figure 2.**
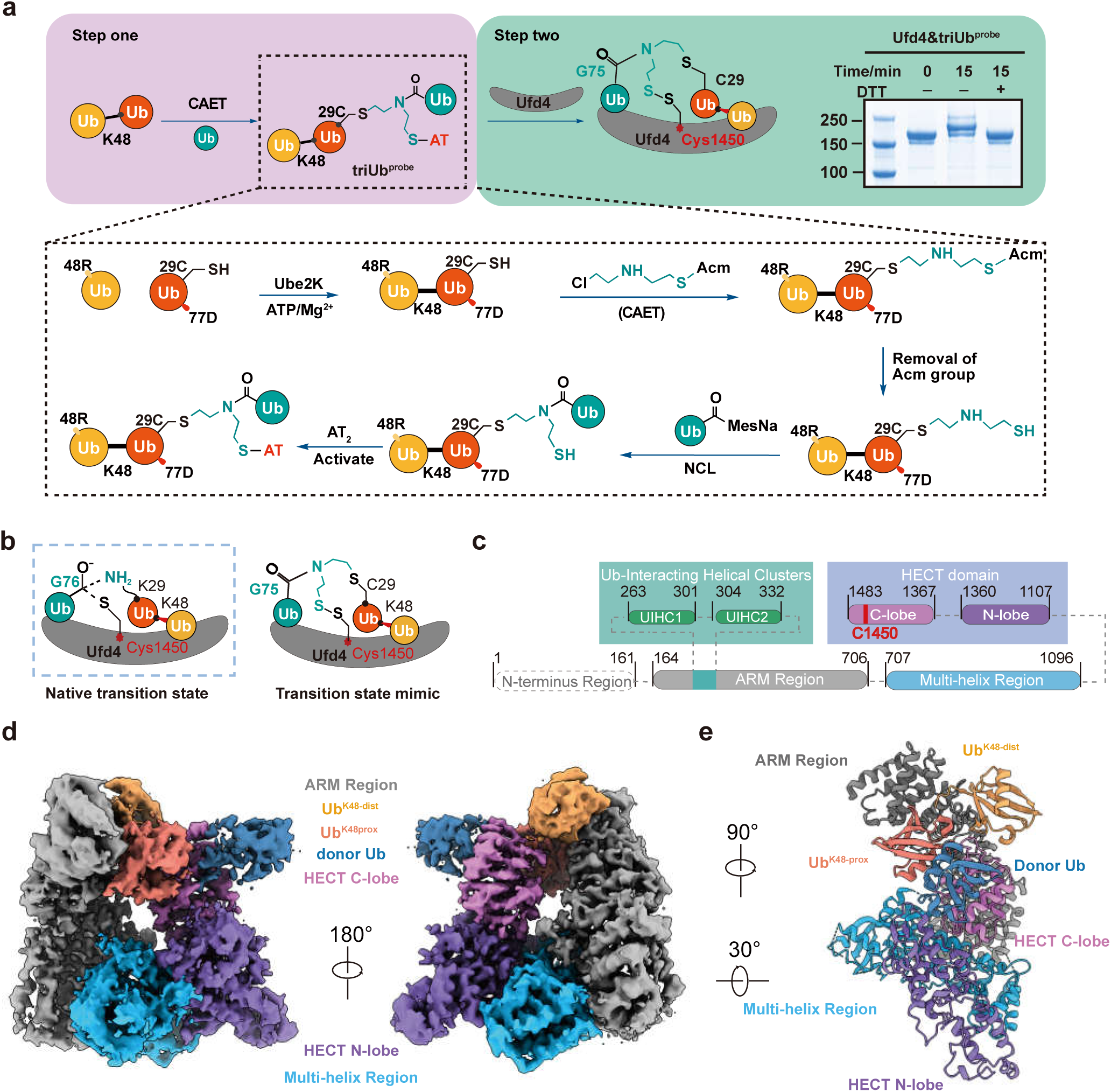
Overall structure of Ufd4 in complex with triUb^K29/K48^. **a**, Schematic of two-step complex assembly routes of Ufd4 in complex with triUb^K29/K48^. Step one: synthetic route of the intermediate structure mimicking the transition state of Ufd4 mediated ubiquitination at proximal Lys29 site of K48-linked diUb. Step two: in vitro crosslinking experiments between Ufd4 and triUb^K29/K48^. **b**, Schematic diagram of the native transition state *vs.* the transition state mimic of the ubiquitination process. The side chain of Lys29 residue on K48-linked diUb attacks the thioester bond of Ufd4–Ub. An intermediate structure design is shown in the inset to mimic the transition state of Ufd4-mediated Ub chain assembly. **c**, Structural domain diagram of Ufd4 with indicated residue boundaries. The dashed-line box indicates unresolved N-terminus region, the turquoise box represents the Ub binding domains, the dusty blue box indicates the HECT domain and the red line represents the active Cys1450. **d, e,** Cryo-EM density (**d**) and structural model (**e**) of the Ufd4-triUb^K29/K48^ complex.

The overall structure of Ufd4 with donor Ub conjugated on proximal K29 of K48-linked diUb (named Ufd4-triUb^K^^29/K48^ complex) resembles a closed ring shape, in which Ufd4 forms a clamp sandwiching donor Ub and K48-linked diUb (Fig. 2d, e). The chemically stable linkage between donor Ub C-terminus, Ufd4 active center, and acceptor Ub K29 mimics the native transition state (Supplementary Fig. 3d). The N-terminal region of Ufd4 (1-163), which was reported to interact with the proteasome^29^, is invisible in our cryo-EM map, possibly due to intrinsic disorder. The visible Ufd4 scaffold consists of three regions: the armadillo-like (ARM) region (164-706) composed of multiple repeating α-helices, the HECT domain that contains the N-lobe (1107-1360) and C-lobe (1367-1483), and the multi-helix region (MHR) (707-1096) bridging the two regions mentioned above (Fig. 2c-e). K48-linked diUb is bound by the C-lobe of the HECT domain and two helical repeats within the ARM region, previously reported to recognize Ub-fusion proteins^30^, which we term here as Ub-interacting helical clusters (UIHC1, residues 263-301; UIHC2, residues 304-332) (Fig. 2c). Meanwhile, the donor Ub is engaged by the C-lobe of the HECT domain, consistent with previous structural findings derived from the HECT domain^13–15^ or full-length UBR5^17^.

### Ufd4 binds the substrate (K48-linked diUb) through multidentate interfaces

The contact between Ufd4 and K48-linked diUb can be subdivided into interactions with proximal Ub (named Ub^K48-prox^) and distal Ub (named Ub^K48-dist^) (Fig. 3a). Ub^K48-prox^ is bound by UIHC2 and the HECT C-lobe, where residues Thr315, Ile316 and His317 of UIHC2 are directed toward residue Arg54 of Ub^K48-prox^ (Fig. 3c and Supplementary Fig. 4f), and residues His1378 and His1433 of the HECT domain C-lobe are pointed toward the hydrophilic surface formed by Ub^K48-prox^ residues Pro19 and Ser20 (Fig. 3d and Supplementary Fig. 4g, 5c). This dual anchoring effect of UIHC2 and the HECT C-lobe on Ub^K48-prox^ orients its Lys29 side chain toward the active cysteine of Ufd4 (distance between α-carbon atoms is 9.04 Å, Supplementary Fig. 5d). Ub^K48-dist^ interacts with the Ufd4 UIHC1 and UIHC2 motifs through multiple hydrophobic interactions. Specifically, residues Tyr276, Ile277 and Asp278 of Ufd4 UIHC1 are located close to the Leu8 loop of Ub^K48-dist^, and the side chain of Phe313 of UIHC2 inserts into the hydrophobic I44 patch of Ub^K48-dist^ formed by Ile44, His68, and Val70 (Fig. 3b and Supplementary Fig. 4e, 5b), which often participate in recruiting Ub-binding proteins^2^. Interestingly, these above interactions reshape K48-linked diUb to adopt an extended conformation that differs from prior structures determined for either free K48-linked diUb or in complex with various recognition proteins (e.g., Rpn13, SARS PLpro, MINDY1)^31–34^ (Supplementary Fig. 5a). Furthermore, the isolated Ufd4 ARM region (residues 164-706) binds to K48-linked diUb with an affinity of 0.47 ± 0.15 μM, while no binding signal was detected in the case of monoUb (Supplementary Fig. 5e). This indicates that the Ufd4 ARM region is a standalone Ub-binding domain for K48-linked diUb, but not for monoUb.

**Figure 3.**
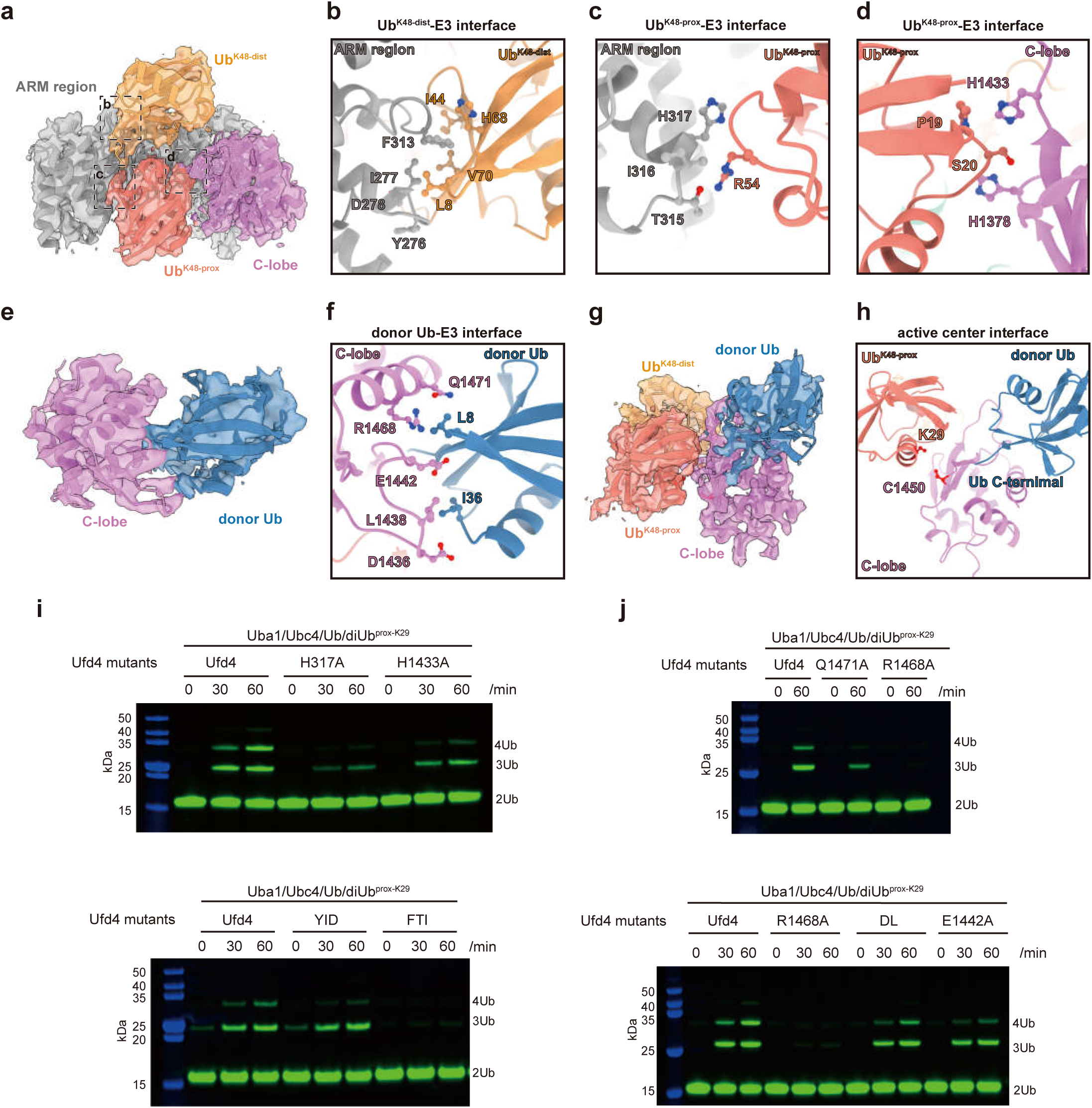
Analysis of the interactions between Ufd4 and triUb^K29/K48^. a, Cryo-EM density of the ARM region and the HECT C-lobe with K48-linked diUb, dashed regions correspond to interfaces between Ufd4 and Ub^K48-dist^, Ufd4 and Ub^K48-prox^. **b**-**d**, Molecular interactions between the ARM region and Ub^K48-dist^ (**b**), the ARM region and Ub^K48-prox^ (**c**), the HECT C-lobe and Ub^K48-prox^ (**d**). **e,** Cryo-EM density of the HECT C-lobe with donor Ub. **f,** Molecular interactions between the HECT C-lobe and donor Ub. **g-h,** Cryo-EM density of the HECT C-lobe and triUb^K29/K48^ (**g**), and the structural model showing the spatial proximity of HECT C-lobe Cys1450, Lys29 of Ub K48-prox and C-terminal of donor Ub (**h**). **i-j**, In vitro Ufd4-dependent ubiquitination assays. Mutants at the substrate recognition interface (**i**) and the donor Ub interface (**j**) were tested. Gel images are representative of independent biological replicates (n = 3). Source data are provided as a Source Data file.

The importance of the above key interacting surfaces was assessed by assaying the effects of mutations in UICH1, UIHC2, and the HECT domain C-lobe of Ufd4. In Ufd4-mediated polyubiquitination of K48-linked diUb, the Ufd4 UIHC1 triple mutant (Y276A/I277A/D278A, named the YID mutant) slows down the activity by 25% (Fig. 3i). The Ufd4 UIHC2 triple mutant (F313A/T315A/I316A, named FTI mutant) or single mutant H317A impairs the activity of Ufd4 by 90% or 70%, respectively (Fig. 3i). The HECT domain C-lobe H1433A mutant reduced activity by a quarter (Fig. 3i). To gain further insight into the specific roles of these residues in branched ubiquitination, we performed enzyme kinetic analysis on the FTI and H317A mutants (Supplementary Fig. 2c, 6a-c), which exhibited the most pronounced defects in ubiquitination activity. Our results showed that, compared to WT Ufd4, both FTI and H317A mutations had minimal impact on the catalytic rate constant (k_cat_), but led to a significant increase in the Michaelis constant (K_m_). This suggests that these residues play a crucial role in substrate binding, rather than in catalysis, which is consistent with their location at the enzyme-substrate interface in the complex.

### Interactions between the Ufd4 HECT domain C-lobe and the donor Ub

The donor Ub is primarily positioned by the C-terminal helix (residues 1462-1475) and the L2 loop (residues 1433-1445) of the C-lobe in the Ufd4 HECT domain (Fig. 3e and Supplementary Fig. 7a). We compared the structural organization between the C-lobe and donor Ub in our Ufd4-triUb^K29/K48^ complex with the canonical arrangement typified by the Ub-loaded NEDD4 HECT domain structure (PDB code: 4BBN)^11^ and E3-Ub-substrate complex (PDB code: 4LCD)^16^ (Supplementary Fig. 7b). Structural alignment based on the C-lobe revealed that the donor Ub in our structures displays a noticeable difference in orientation, exemplified by residue E34 of the donor Ub in our structure flips 40.3° (compared to 4BBN) and 42.8° (compared to 4LCD) relative to C-lobe residue Q1471 (measured by the α carbon atoms) and undergoes shifts of 15.66 Å (compared to 4BBN) and 15.97 Å (compared to 4LCD), respectively.

In the C-terminal helix (1462-1475), residues R1468 and Q1471 are oriented towards the residues L8 and T9 of donor Ub. Similarly, the corresponding C-terminal helix, such as that of NEDD4, HUWE1, Rsp5 and UBR5, has also been observed to interact with donor Ub. Regarding the L2 loop (1433-1445), it contains an insertion of five amino acids compared to the homologous loops, which makes itself reach out and guide the rotation of donor Ub. Residues D1436, L1438, and E1442 are proximal to the hydrophobic patch of donor Ub formed by residues I36 and L71 (Fig. 3f and Supplementary Fig. 4h), and notably, residues D1436 and L1438 are located within this insertion. Consistent with these structural observations, the Ufd4 R1468A mutant significantly impaired both Ufd4∼Ub thioester formation and ubiquitination of the K48-linked diUb substrate, while the Ufd4 E1442A mutant and the double mutant (D1436A/L1438A) exhibited varying reductions in Ufd4∼Ub thioester formation and substrate ubiquitination activity (Fig. 3j and Supplementary Fig. 7c, d). The above interaction positions the C-terminal thioester of the donor Ub proximal to the K29 side chain of Ub^K48-prox^, underlying the structural insight into Ufd4-catalyzed K29-linked branched ubiquitination on K48-linked Ub chain (Fig. 3g, h).

### Validation of key interactions between Ufd4 and K29/K48-branched triUb in vivo

To investigate the effect of the key interfaces observed in our structure, a 43-mer type-1 degron peptide and a truncated substrate Scc1 (residues 268-384) were used in the Ufd4-mediated polyubiquitination assay. The K48-linked Ub chains were first preassembled on these two substrates using Ubr1 and its E2 enzyme Ubc2, and then further elongated by Ufd4 and its mutants. The ubiquitination results showed that WT Ufd4 could further extend the polyubiquitin chain on the type-1 degron peptide as well as the bona fide substrate Scc1 (residues 268-384), while the Ub chain elongating activity was substantially reduced by the Ufd4 mutants FTI, R1468A, and H317A (Supplementary Fig. 8a, b, f).

To validate the structural mechanism in a physiological setting, a previously developed yeast growth assay was performed^19^. In *S. cerevisiae*, the ubiquitination level of Mgt1 is regulated by both Ufd4 and Ubr1, and its degradation leads to yeast death when exposed to the DNA alkylating agent MNNG^19^. We constructed a Ufd4-knockout yeast strain, and complementation with Ufd4 or its mutants altered Mgt1 ubiquitination levels, thus affecting the growth status of yeast (Supplementary Fig. 8c). When WT Ufd4 was introduced to complement the Ufd4-knockout yeast strain, exposure to the DNA-alkylating agent MNNG impaired the growth of the complemented strain. In contrast, under conditions without MNNG treatment, the growth phenotype of the Ufd4-knockout strain closely resembles that of the strain complemented with WT Ufd4 (Supplementary Fig. 8d). These results were consistent with previous in vivo degradation experiments showing that the Mgt1 level was affected by Ufd4^19^. Complementation with Ufd4 mutants (FTI, R1468A, or H317A) decelerated yeast death compared to WT Ufd4 with the addition of MNNG (Supplementary Fig. 8d). Moreover, the Ufd4-knockout strain was transformed with Myc-tagged Mgt1 and wild-type Ufd4 (UFD4^WT^) or different mutants (UFD4^FTI^, UFD4^H317A^ and UFD4^R1468A^), followed by a chase for 10 min to evaluate the decrease in Mgt1 protein levels mediated by different Ufd4 mutants. Complementation with UFD4^WT^ was observed to promote the degradation of Mgt1, whereas complementing with mutated UFD4 reduced the degradation efficiency of Mgt1 (Supplementary Fig. 8e). These results support the correlation between the key interfaces of Ufd4 with K29/K48-branched triUb and their functional effects, highlighting the crucial role of these interactions in regulating Mgt1 degradation.

### Conserved mechanism of K29/K48-branched ubiquitin chain formation by TRIP12

TRIP12 (the human homologue of Ufd4) has been discovered to be a positive regulator of PROTAC-induced neo-substrate degradation, facilitating this process by cooperating with the K48-linked Ub chain-specific CRL2 complex to form K29/K48-branched Ub chains^10^. This finding parallels Ufd4’s preference for K48-linked Ub chains in generating K29/K48-branched Ub chains. We reconstructed the TRIP12-catalyzed ubiquitination using K48-linked diUb as a substrate and observed polyubiquitination of the substrate. Similar to Ufd4, TRIP12 exhibits the strongest ubiquitination activity towards K48-linked diUb substrates among the eight different diUb linkage types (Supplementary Fig. 9a, b). Middle-down MS analysis of the polyubiquitination product revealed 32.3% of mono-Ub species modified by double-glycine remnants (Supplementary Fig. 9d), representing the generation of branched Ub chains by TRIP12. Additionally, Ub fragments with double-glycine remnants on both K29 and K48 residues were detected in the MS/MS spectrum (Supplementary Fig. 9e). Furthermore, we performed ubiquitination assays using K48-linked diUb mutants with K29R in either the proximal or distal Ub as substrates. The biochemical results demonstrated that mutants with proximal K29R mutations exhibited weak ubiquitination activity, whereas mutants with distal K29R mutations remained efficiently ubiquitinated (Supplementary Fig. 9c), supporting the preference of TRIP12 for the K29 site on proximal Ub to generate K29/K48-branched Ub chains. These results suggest that TRIP12, like Ufd4, prefers K48-linked Ub chain substrates to assemble K29/K48-branched Ub chains.

To visualize the catalytic assembly of K29/K48-branched Ub chains by TRIP12, triUb^probe^ was also employed to obtain catalytic complex mimics, similar to the approach utilized for Ufd4. We determined a cryo-EM structure of the TRIP12/triUb^probe^ complex at 3.63 Å, facilitated by the AlphaFold2-predicted TRIP12 structure and the crystal structure of Ub as an initial template for modelling (Supplementary Tab. 1 and Supplementary Fig. 22). For TRIP12, the armadillo-like (ARM) region (442-1106), the multi-helix region (MHR) (1107-1584), and the catalytic HECT domain were docked unambiguously (Fig. 4a, b). Among the three Ub units, the map was best defined for the substrate K48-linked diUb. However, the map density for donor Ub was weak. The active Cys of TRIP12 and the acceptor Ub residue K29 are located in the vicinity of 6.52 Å.

**Figure 4.**
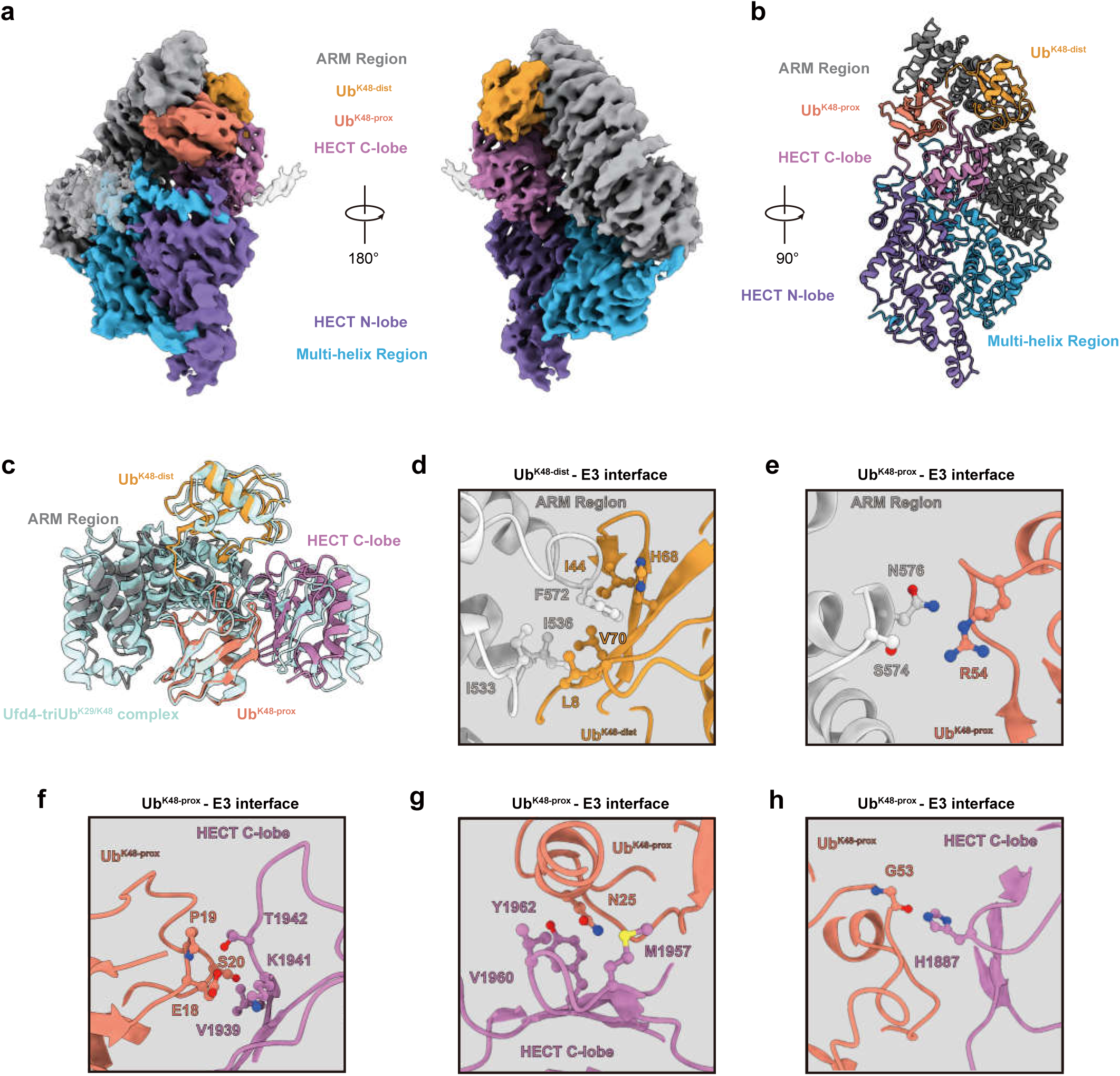
Analysis of the interactions between TRIP12 and triUb^K29/K48^. a, Cryo-EM density (**a**) and structural model (**b**) of the TRIP12-triUb^K29/K48^ complex. **c**, Structural alignment of Ufd4-triUb^K29/K48^ complex and TRIP12-triUb^K29/K48^ complex structures, and only ARM region, C-lobe and K48-linked diUb are shown. **d**-**e**, Molecular interactions between the ARM region and Ub^K48-dist^ (**d**), the ARM region and Ub^K48-prox^ (**e**). **f**-**h**, Molecular interactions between the C-lobe and Ub^K48-prox^.

In the complex, TRIP12 binds to the substrate K48-linked diUb in a shape similar to that of the Ufd4/triUb^K29/K48^ complex, with the structural organization of the substrate remaining virtually unchanged (Fig. 4c). The ARM region and the C-lobe of the HECT domain synergistically clamp the K48-linked diUb, mirroring the recognition framework of Ufd4 (Supplementary Fig. 9f). The TRIP12 fragments corresponding to UICH1 and UICH2 interact with the distal Ub mainly through hydrophobic interactions, and the interaction network closely resembles that observed in the Ufd4-distal Ub interaction. Specifically, the L8 loop of distal Ub interacts with TRIP12 residues I533, I536, and F572. Meanwhile, the I44 patch in distal Ub binds to residues F572 and I536 of TRIP12 (Fig. 4d). These key residues of TRIP12 involved in interactions with the distal Ub are conserved from yeast to humans, implying an important role for this interface in Ub recognition. For proximal Ub, as in the

Ufd4/triUb^K29/K48^ complex, it engaged with UICH2 and the C-lobe. The proximal Ub residues R54 and P19/S20, which also participate in binding with Ufd4 in the Ufd4/triUb^K29/K48^ complex, interact with residues S574/N576 in UICH2 and T1942/V1939 in the C-lobe of TRIP12, respectively (Fig. 4e, f). Compared to the Ufd4/triUb ^K29/K48^ complex, the C-lobe of TRIP12 undergoes a slight shift, positioning it closer to the proximal Ub (Fig. 4c). The cryo-EM density map indicated that E18 of the proximal Ub establishes an ionic interaction with K1941 of the C-lobe (Fig. 4f); N25 in the proximal Ub inserts into the hydrophobic cavity formed by M1957, V1960, and Y1962 on the C-lobe (Fig. 4g); and the backbone of G53 is adjacent to H1887 of the C-lobe, facilitating backbone hydrogen bonding interaction (Fig. 4h). Although these interactions were not observed in the Ufd4/triUb ^K29/K48^ complex due to species differences, they did not reshape the proximal Ub orientation, which remains the same as that observed in the Ufd4/triUb complex. To further explore the functional significance of these key interactions, we generated TRIP12 mutants, including I533G/I536G/F572G, V1939G/K1941A/T1942G, and M1957G/V1960G/Y1962G, and subjected them to ubiquitination assays. The biochemical results showed that these mutations significantly reduced ubiquitination activity (Supplementary Fig. 9g), emphasizing the crucial role of these interactions in TRIP12’s catalysis of K29/K48-branched Ub chain formation. Collectively, our biochemical and structural analysis revealed a conserved enzymatic property and structural mechanism to form the K29/K48-branched Ub chain by Ufd4 and TRIP12, providing insights into how TRIP12 accelerates PROTAC-induced ubiquitination degradation of neo-substrates.

### Cryo-EM structure of Ufd4 in complex with E2 enzyme Ubc4

To obtain a detailed understanding of the catalytic cascade of Ufd4, we determined the structure of Ufd4 accepting Ub thioester from Ubc4. Thus, the active site of Ufd4, the C-terminus of Ub and the active site of Ubc4 were chemically linked by a two-step method similar to that described above (Fig. 5). An activity-based probe that relies on a chemically engineered Ubc4-Ub (named Ubc4-Ub^probe^) was first synthesized and then crosslinked with Ufd4 in an active center (C1450)-dependent manner to form a stable complex mimic (Fig. 5a, b and Supplementary Fig. 3b, c). The mimic was subjected to cryo-EM analysis and provided three cryo-EM reconstructions, including a 3.52 Å map in which Ub and Ufd4 (residue 1-706) were invisible, a 4.30 Å map with Ub and Ufd4 (residue 1-161) were invisible, and a 6.39 Å map showing Ufd4 (residue, 1-706) was invisible while Ub was visible (Supplementary Tab. 1 and Supplementary Fig. 23).

**Figure 5.**
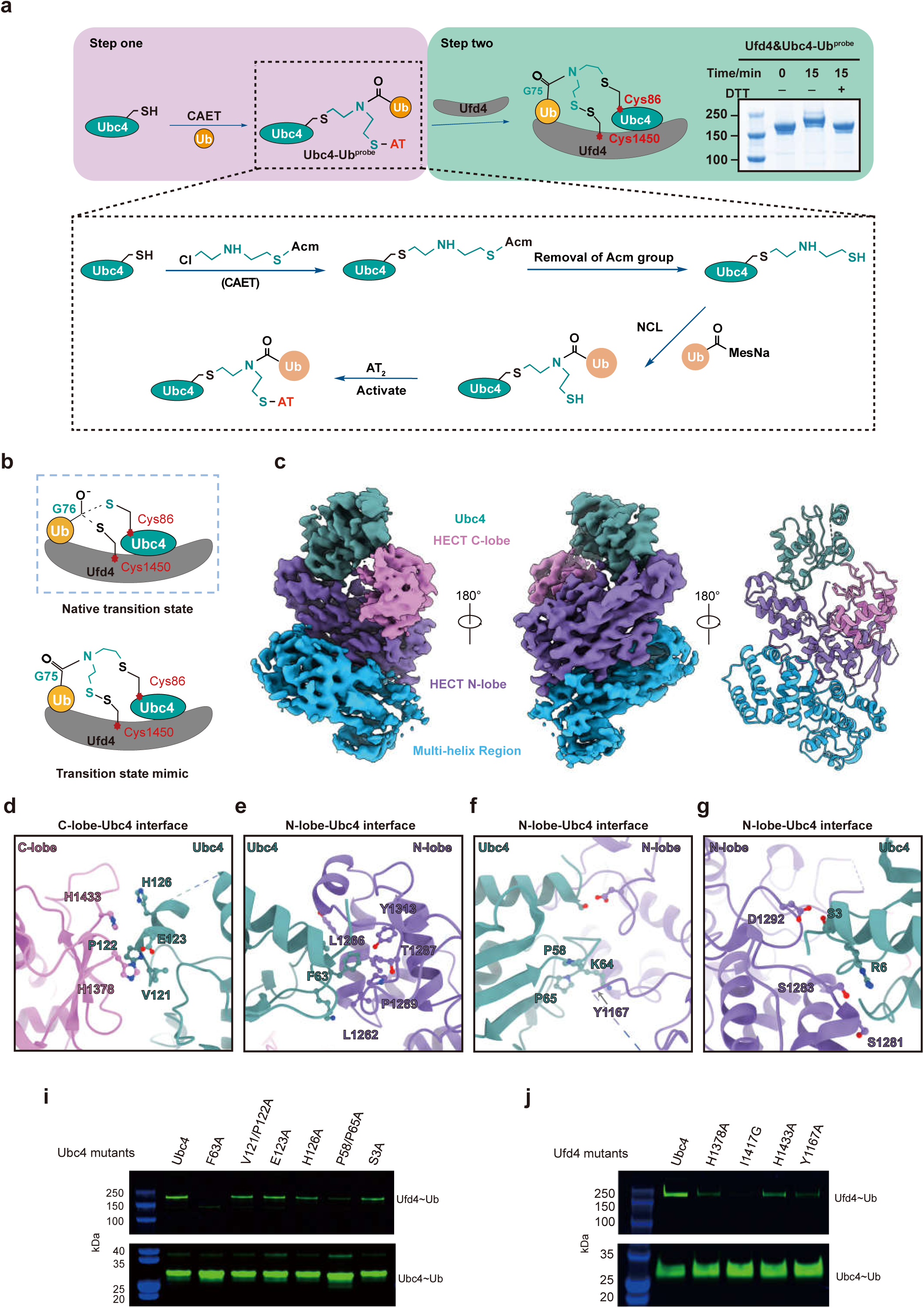
The structure of Ufd4 in complex with Ubc4-Ub. a, Schematic of two-step complex assembly routes of Ufd4 in complex with Ubc4-Ub. Step one: synthetic route of the intermediate structure mimicking the transition state of Ufd4-mediated transthiolation from Ubc4-Ub. Step two: in vitro crosslinking experiments between Ufd4 and Ubc4-Ub. **b**, Schematic diagram of the native transition state vs. the transition state mimic of the transthiolation process. The side chain of catalytic Cys residue on Ufd4 attacks the thioester bond of Ubc4–Ub. An intermediate structure design is shown in the inset to mimic the transition state of the Ufd4-mediated transthiolation process. **c**, Cryo-EM density and structural model of the Ufd4^△ARM^-Ubc4 complex. **d**-**g**, Molecular interactions between the HECT C-lobe and Ubc4 (**d**), the HECT N-lobe and Ubc4 (**e**-**g**). **i-j**, In vitro Ufd4∼Ub thioester formation assays. Ubc4 mutants (**i**) and Ufd4 mutants (**j**) were tested. Gel images are representative of independent biological replicates (n = 2). Source data are provided as a Source Data file.

The primary distinction among the three maps above lies in the visibility of Ub and the N-terminal region of Ufd4, and across all three maps, there is generally good overlap in the density of the Ufd4 HECT domain and Ubc4 (Supplementary Fig. 10a). Detailed structural analysis of Ufd4 with Ubc4 was then conducted by focusing on the highest resolution structure (3.52 Å), facilitated by the AlphaFold-predicted Ufd4 structure (code: AF-P33202-F1) and the crystal structure of Ubc4 (PDB code: 1QCQ) as a starting point for model building (Supplementary Fig. 4c, d). In this structure, the catalytic cysteines of Ubc4 and Ufd4 are positioned closely, with an α-carbon distance of less than 8 Å, suggesting that this structure may represent an active trans-thioesterification state, though the Ub is not visible.

Ubc4 is embraced by both the N-lobe and C-lobe of the Ufd4 HECT domain, exhibiting a similar arrangement to the canonical HECT E3-E2 transthiolation complex, as exemplified by the NEDD4L-Ubch5b-Ub complex^11^ (Fig. 5c, Supplementary Fig. 10b). The contacts between the Ufd4 HECT domain N-lobe and Ubc4 are contributed by multiple interfaces. The residue F63 of Ubc4, crucial for determining the specificity of E2/E3 interaction, fits snugly into a hydrophobic groove formed by L1262, L1266, I1269, L1284, Y1313, and other residues (Fig. 5e and Supplementary Fig. 4j), reminiscent of how the homologous Phe residue in Ubch7 or Ubch5b binds to the parallel hydrophobic surface of the E6AP or NEDD4L HECT domain. Indeed, the Ubc4 F63A mutant almost completely diminished the trans-thioesterification activity (Fig. 5i). Three other interactions between N lobe and Ubc4 are presented: the N-lobe residue D1292 and Ubc4 residue S3; the N-lobe residue S1281/S1283 and Ubc4 residue R6; the N-lobe residue Y1167 and Ubc4 residues P58, K64 and P65 (Fig. 5f, g and Supplementary Fig. 4k, l). Mutations in these interface residues are deleterious for the formation of E3∼Ub thioester (Fig. 5i, j). The interaction between the Ufd4 HECT domain C-lobe and Ubc4 involves the pairing of C-lobe residue H1433 with Ubc4 residue H126 and the pairing of C-lobe residue H1378 with Ubc4 residues V121, P122, and E123 (Fig. 5d and Supplementary Fig. 4i). The Ufd4 mutants H1433A and H1378A, as well as the Ubc4 mutant E123A and double mutant V121A/P122A, all reduced the transfer of Ub from Ubc4 to Ufd4 (Fig. 5i, j).

In our structure, the HECT domain N- and C-lobes adopt the inverted T-conformation, as observed in the NEDD4L and UBR5 transthiolation complex^11,17^. The interaction between N lobe residues R1224, K1152, Y1181 and T1183, and residue I1417 on the C-lobe, plays a potential role in stabilizing this conformation (Supplementary Fig. 11b). Indeed, when residue Ile1417 was mutated to Gly1417, the E3∼Ub thioester formation efficiency was reduced to ∼20% compared to WT Ufd4 (Fig. 5j) and the branched ubiquitination was significantly impaired (Supplementary Fig. 10c). In conclusion, Ubc4 and Ufd4 coordinate Ub transfer through a module similar to that described previously^11,12^.

### Transition model of Ufd4 from transthiolation to K29-linked Ub assembly

The two structures we have resolved above correspond to the HECT E3 enzyme E2-to-E3 transthiolation state (Ufd4-Ubc4 complex structure) and the E3-to-substrate ubiquitination state (Ufd4-triUb^K29/K48^ complex structure). The similarities and differences between these two conformations were analyzed to investigate the interdomain conformational changes of the HECT E3 enzyme in different states.

First, we focused on the conformational differences of Ufd4 in complex with Ubc4 and triUb. The structures of Ufd4-Ubc4 (which contains the ARM region) and Ufd4-triUb^K29/K48^ were compared, and the Ufd4 MHR (707-1096) and the N-lobe (1361-1107) superimpose well in these two structures (Supplementary Fig. 11a). Notably, the ARM region (163-706) in the two structures undergoes conformational changes; for instance, the Ufd4 residue Glu184 undergoes a 28.41 Å shift and a 14.8° deflection relative to the residue His715. Next, we compared the structures of Ufd4-Ubc4 (the structure at the highest resolution) and Ufd4-triUb^K29/K48^ to identify the conformational changes in the HECT domain of Ufd4 in different catalytic states. The structural organization of the HECT domain N-lobe in these two structures remains unchanged during state transitions (Supplementary Fig. 11c), whereas, similar to ubiquitination catalyzed by Rsp5 and UBR5, the C-lobe of the HECT domain adopts distinct conformations during two different reaction states (transthiolation and substrate ubiquitination) (Supplementary Fig. 11c). Finally, the Ufd4 C-lobe residues H1378 and H1433 were found to interact with both Ubc4 and K48-linked diUb in the Ufd4 -Ubc4 and Ufd4-triUb^K29/K48^ structures, respectively (Fig. 3d, 5d), indicating that Ubc4 may need to disassociate beforehand and the C-lobe reorientates to provide space for interactions with K48-linked diUb during the ubiquitination process.

Based on the structures of Ufd4-mediated Ub charging and ubiquitin transfer to the proximal Lys29 site of K48-linked diUb, we proposed a functional model representing the whole catalytic process of Ufd4. First, Ubc4∼Ub is recruited by the HECT domain of Ufd4, followed by the loading of Ub from Ubc4 and the discharge of Ubc4. Subsequently, the conformation of the ARM region and C-lobe reorientates to jointly sandwich the K48-linked diUb substrate, which directs the proximal Lys29 residue of K48-linked diUb toward Ufd4 catalytic Cys1450. Finally, the Ub thioester transfers from Ufd4 catalytic Cys1450 to the proximal Lys29 of K48-linked diUb (Supplementary Fig. 11d).

## Discussion

Prior to proteasomal degradation, substrate proteins with destabilizing residues at the N-terminus (termed as N-degrons) are often recognized by UBR family E3 ligases and ubiquitinated in a K48 linkage-specific manner^35^. This ubiquitination process on N-degron substrates can be further augmented by another E3 ligase Ufd4 (a K29-linkage-specific HECT-type E3 ligase), leading to accelerated substrate degradation^9,19^. The mechanism of a representative yeast Ubr1-catalyzed process, including the initiation of N-degron ubiquitination and the K48 linkage-specific elongation step, was recently elucidated by cryo-EM studies in our group^36^. Nonetheless, the mechanism through which Ufd4 introduces K29-linked Ub chains on substrates to increase the processivity of polyubiquitination remains unknown. Here, our biochemical and mass spectrometry experiments clarified key aspects of the molecular mechanism of Ufd4, where Ufd4 preferentially catalyses branched ubiquitination on K48-linked Ub chains, thereby elongating the Ub chain. To gain structural insight into this process, we designed a mechanism-based chemical trapping strategy to study the Ufd4-specific ubiquitination processes, specifically the transfer of Ub from the E2 enzyme to the E3 ligase, and subsequently from the E3 ligase to the substrate. This strategy combined with cryo-EM provides valuable mechanistic insights into how Ufd4 mediate the K29/K48-branched ubiquitination and enhances our understanding of the catalytic mechanism underlying HECT E3 ligases. Meanwhile, our findings contribute to a deeper understanding of linkage-specific polyubiquitin chain assembly by providing mechanistic details specific to K29-linked ubiquitination, thereby complementing previous structural studies on M1-, K11-, K48-, and K63-linked Ub chains^17,36–42^. Interestingly, the human homologue of Ufd4, TRIP12, also preferentially assembles K29/K48-branched Ub chains through a conserved structural mechanism, suggesting an evolutionarily conserved function across species.

A variety of structural elements responsible for acceptor Ub recognition have been identified within the catalytic domains of E2 and E3 enzymes, which play a critical role in assembling Ub chain^17,36–42^. For instance, a flexible loop in the E2 enzyme UBE2R2 is reshaped by the CRL RING domain to precisely position the acceptor Ub within the catalytic site^42^. Likewise, a zinc finger motifembedded in the RING2 domain of the RBR-type E3 ligase RNF216 engages the acceptor Ub, thereby ensuring the selective generation of K63-linked Ub chains^40^. Despite these insights, the contribution of non-catalytic regions in E2 and E3 enzymes to acceptor Ub recognition remains poorly understood. A recent structural study on the HECT-type E3 ligase UBR5, which specifically assembles K48-linked Ub chains, revealed that its N-terminal UBA domain binds an acceptor Ub, emphasizing the crucial role of non-catalytic domains in dictating Ub chain assembly^17^. In this study, we found that the non-catalytic N-terminal ARM domain of Ufd4/TRIP12 cooperates with the HECT catalytic domain to precisely position the acceptor Ub, aligning its Lys29 toward the Ufd4/TRIP12 active site. Although the N-terminal ARM domain may play a secondary role in this process, its involvement further underscores the emerging concept that non-catalytic regions in Ub ligases contribute to Ub chain assembly. Moving forward, it will be essential to investigate Ub chain assembly at the level of full-length ubiquitination enzymes rather than focusing solely on their catalytic domains. Such studies will provide deeper insights into the role of non-catalytic domains in acceptor Ub recognition.

Currently, structural studies on homotypic Ub chain assembly primarily focus on the formation of diUb chains, where only the acceptor Ub unit needs to be recognized for chain elongation. In contrast, a structural model for the generation of longer homotypic Ub chains has been established mostly through the structural study of the UBE2K/RNF38 RING domain, which catalyzes the formation of K48-linked triUb from K48-linked diUb^39^. In this model, only the acceptor (distal) Ub is recognized, while the proximal Ub remains unbound. This raises the intriguing question of whether branched Ub chain formation follows a similar recognition pattern. Specifically, in the case of branched ubiquitination on a diUb substrate, does the recognition of the proximal Ub alone suffice to initiate the branching process? Here, we found that for Ufd4/TRIP12, when K48-linked diUb serves as the substrate, both the proximal and distal Ubs are recognized simultaneously. This suggests that, at least in the case of Ufd4/TRIP12, unlike homotypic chain elongation, which relies solely on the recognition of the distal Ub, the formation of branched Ub chains requires the recognition of multiple Ubs. Building on this, we developed a working model for branched Ub chain assembly, highlighting the key distinctions between homotypic and branched Ub chain formation. This observation raises the question of whether the recognition of multiple Ubs for the position of the acceptor Ub is a common feature among E3 ligases catalyzing branched ubiquitination, or if it is unique to Ufd4/TRIP12. We anticipate that future structural studies on other branched Ub ligases will provide further insights.

## Methods

### Cloning and plasmid construction

The plasmid encoding *S. cerevisiae Ufd4* gene was synthesized by GenScript (Nanjing, China), and its codon was optimized for *S. cerevisiae* and *Escherichia coli* (*E. coli*) overexpression, respectively. For overexpression by *S. cerevisiae*, the *Ufd4* gene was subcloned to the vector YEplac181 containing an N-terminal DYKDDDDK (Flag) tag. For overexpression by *E. coli*, the gene was inserted between the BamHI and EcoRI sites of the vector pGEX-4T-1 with an N-terminal GST tag followed by an HRV 3C protease cleavage site. The plasmid encoding human TRIP12 (442-1992) was also synthesized by GenScript, and its codon was optimized for HEK293F overexpression, which was subcloned to the vector pcDNA3.1 containing an N-terminal Flag tag. The DNA sequence encoding yeast (*S. cerevisiae*) *Ubc4* was synthesized by GenScript and its codon was optimized for *E. coli* overexpression. The gene was further cloned into the vector pET-28a with an N-terminal His tag followed by an HRV 3C protease cleavage site. Different *Ufd4* and *Ubc4* mutants were constructed by site-directed mutagenesis. The DNA sequences of wild-type Ub and Ub mutants including Ub-K6R, Ub-K11R, Ub-K27R, Ub-K29R, Ub-K33R, Ub-K48R, Ub-K63R, Ub-G76C, Ub-K29R-G76C, Ub-K29R/K48R, Ub-K29C/K48R, Ub-K29R/K63R, Ub-77D (a Ub variant with an additional amino acid Asp at the C terminus), Ub-K29C/77D, M1-linked triUb mutant (distal and intermediate ubiquitin with K29R mutation, proximal ubiquitin with G76C mutation) and Ub-MCQ (Ub variant with an additional amino acid Cys between Met1 and Gln2) were constructed and cloned to the vector pET-22b by GenScript. The plasmids containing *S. cerevisiae/*human *Uba1*, *S. cerevisiae Ubc2/*human *Ubch7*, *S. cerevisiae Ubr1*, and the truncated *S. cerevisiae* genes *Scc1* (268–384; also known as MCD1) were the same as previously used^12,36,43,44^.

### Protein expression and purification

Ufd4 used for cryo-EM sample and its mutants used in Ufd4-triUb^K29/K48^ complex structure-related ubiquitination assay were expressed in yeast and purified similarly to yeast Ubr1 as previously described^36^. In brief, plasmids containing *Ufd4* and its mutants were transformed into yeast (BY4741) cells. Single colonies were selected to grow in a synthetic medium with Leu defective resistance (SC, −Leu) to an optical density at 600 nm (OD_600_) of 2-3. The yeast cells were collected by centrifugation at 5,000 × *g* and then fully resuspended with lysis buffer (50 mM HEPES, pH 7.5, 150 mM NaCl, protease inhibitor cocktail (Roche), and 10% glycerol). The suspension liquid was frozen into small balls and subsequently ground into a fine powder using a freezing grinder (SPEX Sample Prep 6870 Freezer/Mill). The powder was defrosted in ice water and centrifuged at 17,418 × *g* for 1 h. The supernatant was loaded on the anti-Flag affinity resin (Thermo Fisher Scientific, A36803) and incubated at 4℃ for 2h. After washing adequately, 1 mg/mL Flag peptide dissolved in buffer (50 mM HEPES, pH 7.5, 150 mM NaCl) was used to elute Flag-tagged Ufd4. Ufd4 was then purified by ion exchange (Mono Q anion exchange column, GE Healthcare) with buffer A containing 50 mM HEPES, pH 8.0 and buffer B containing 50 mM HEPES, pH 7.5, 1 M NaCl. Samples at indicated peaks were concentrated and further purified using size exclusion chromatography (Superose 6 column, GE Healthcare) with a buffer containing 50 mM HEPES, pH 7.5, and 150 mM NaCl. Finally, Ufd4 was analyzed by SDS-PAGE (LabPAGE 4-12% 15 Wells, LABLEAD). Human TRIP12 (442-1992) was expressed using HEK293F (Expi293F) cells and initially purified using anti-Flag affinity resin and further purified using ion exchange (Mono Q) and exclusion chromatography (Superose 6 column), similar to that of Ufd4.

The purification of Ubc2, Ubc4 and its variants were the same as previously^36^ described. Briefly, plasmids containing *Ubc4* and its variants were introduced into *E. coli* BL21(DE3) competent cells and cultured in Luria Broth (LB) medium with 20 μg/mL kanamycin at 37℃ to OD_600_ of 0.6. The cells were cooled to 18℃ and induced by 0.4 mM isopropylβ-d-1-thiogalactopyranoside (IPTG) for 16h. The cells were centrifuged at 5,000 × *g* for 25 min at 4 °C and the precipitate was fully resuspended with lysis buffer (50 mM HEPES, pH 7.5, 150 mM NaCl and 1 mM phenylmethyl sulfonyl fluoride (PMSF). After lysed by sonication and 17,418 × *g* centrifugation, the supernatant was incubated with Ni-NTA affinity column for 2 h at 4 °C. After washing with at least 10 column volumes of lysis buffer (50 mM HEPES, pH 7.5, 150 mM NaCl, 20 mM imidazole), the proteins were eluted with a lysis buffer containing 400 mM imidazole and further purified with size exclusion chromatography (Superdex 200 column, GE Healthcare) with a buffer containing 50 mM HEPES, pH 7.5, and 150 mM NaCl. The expression of Ufd4 used in the Ufd4/Ubc4-Ub complex structure-related ubiquitination assay was similar to Ubc4 as we described above. Differently, after lysis by sonication and high-speed centrifugation, the supernatant was incubated with Glutathione beads for 2 h at 4℃. The protein was eluted with a lysis buffer containing 30 mM Glutathione. The elution was incubated with HRV 3C protease overnight to remove the GST tag and further purified as described above for yeast-expressed Ufd4. Scc1(268–384) and Uba1 were obtained as we used before^36^. Wild-type (WT) Ub and Ub mutants were purified as previously described^36^.

### Yeast growth assay

*S. cerevisiae* strain lacking *Ufd4* (*ΔUfd4*) was constructed by Thermo Fisher Scientific (item number: YKL010C_34859). Ufd4 mutants (R1468A, FTI mutant, H317A) were constructed in the context of the vector YEplac181, carrying the wild-type Ufd4.

Yeast strains lacking Ufd4 were plated in a rich (YPD) solid medium containing 2% agarose and 2% glucose (w/v) and grown at 30℃ for 2 days. Single colonies were selected to grow in a YPD liquid medium at 30℃ to mid-log-phase (OD_600_ of 0.4-0.8). Yeast cultures were centrifuged at 1,962 × *g* for 10 min and washed with double-distilled H_2_O. Centrifuged again and resuspended with 1×TE buffer (10 mM Tris-HCl, pH 8.0, 1 mM Ethylene Diamine Tetraacetic Acid (EDTA)). Repeated and resuspended with 1×LiAc (100 mM LiAc, pH 7.5) and 0.5×TE buffer to obtain *ΔUfd4* competent yeast cells and stored on ice for further use. WT Ufd4 and its mutants (R1468A, FTI mutant, H317A) were transformed to *ΔUfd4* competent cells and plated in SD (−Leu) medium containing 2% agarose and 2% glucose (w/v) at 30℃ for 2 days.

Single colonies of the yeast strains mentioned above were selected to grow in SD (−Leu) medium containing 2% glucose at 30℃ overnight. Dilute with SD (−Leu) medium to OD_600_ of 0.4 and grow at 30°C until OD_600_ of 1.0. Add N-methyl-N’-nitro-N-nitrosoguanidine (MNNG) stock solution (50 mM NaOAc, pH 5.0, 6 mM MNNG) to the final concentration of 60 μM and grow at 30°C for 1 h As previously described^19^. Single colonies were spotted in serial fivefold dilution onto the SD (−Leu) plates. The plates were grown at 30°C for 2 days before imaging.

### Preparation of Ub chains

K29-linked chains were obtained with 0.2 μM Uba1, 4 μM Ubc4, 0.5 μM Ufd4, and 200 μM WT Ub at 30°C in the reaction buffer (50 mM HEPES, pH 7.5, 150 mM NaCl, 10 mM MgCl_2_ and 5 mM ATP). K48-linked chains were obtained with 1 μM Uba1, 20 μM UBE2K and 800 μM WT Ub at 30°C in the same reaction buffer. For K48-linked diUb variants, WT Ub was substituted with Ub mutants: diUb^distal-K^^29^(200 μM Ub-K29R/G76C and 200 μM Ub-K48R); diUb^prox-K29^(200 μM Ub-G76C and 200 μM Ub-K29R/K48R); diUb^K29C^(200 μM Ub-K29C/77D and 200 Ub-K48R). K48diUb was used for kinetic assay (200 μM Ub-G76C and 200 μM Ub-K48R). The reactions were terminated by buffer (50 mM NaOAc, pH 4.5) and K29-linked and K48-linked chains were separated by ion exchange (Mono S cation column, GE Healthcare) with buffer A containing 50 mM NaOAc, pH 4.5 and buffer B containing 50 mM NaOAc, pH 4.5, 1 M NaCl. Samples at indicated peaks were collected and then dialyzed into a buffer containing 50 mM HEPES, pH 7.5 and 150 mM NaCl, for further use.

For the triUb used in Figure 1f, E2 enzymes and deubiquitinating enzyme (YUH1) were combined to prepare them (Supplementary Chemistry File 9). Briefly, Uba1(1μM), UBE2K or UBE2V/UBE2N (20 μM) and corresponding ubiquitin mutants (800 μM each) were incubated in the reaction buffer at 37°C for 5-8 hours to generate the diUb variants. 7‰ trifluoroacetic acid (v/v) was added to terminate the reaction, then the supernatant was collected and pH adjusted to 7.4, followed by the addition of YUH1 (0.8 μM) for overnight reaction. The diUb products were purified by ion exchange (Mono S cation column, GE Healthcare) and dialyzed into a buffer containing 50 mM HEPES, pH 7.4 and 150 mM NaCl, for further reaction. Next, the above diUb (400 μM), Ub-G76C or Ub-K29R-G76C (400 μM), Uba1(1 μM), UBE2K or UBE2V/UBE2N (20 μM) were mixed and reacted at 37°C for 5-8 hours to generate the triUb variants. and purified as above. M1/K6/K27/K29/K33/K48/K63-linked diUb were obtained as we used before^44^.

### Preparation of fluorescently labeled substrates

Ub-MCQ, Ub-G76C, diUb^prox-K29^, diUb^distal-K29^, K48diUb used for the kinetic assay, triUb used in Figure 1f were obtained as described above and were subjected to label. For the labeling reaction, 1.5 eq. of fluorescein-5-maleimide (Invitrogen, F150) was added to 2 mg Ub or Ub chain mutants, and the mixtures were incubated at room temperature for 2 hours. The labeled proteins were purified with size-exclusion chromatography (Superdex 200 column, GE Healthcare) with a buffer containing 50 mM HEPES, pH 7.5, 150 mM NaCl.

### *In vitro* Ubiquitination assay

*In vitro* ubiquitination assays were performed with 0.2 μM Uba1, 4 μM Ubc4 or Ubch7, 0.25 μM Ufd4 or TRIP12, 10 μM substrate Ub chains (M1/K6/K27/K29/K33/48/K63-linked diUb, K48-linked diUb/triUb/tetraUb/pentaUb) or 2 μM fluorescently labeled substrates (fluorescent Ub-G76C, K48-diUb, diUb^prox-K29^, diUb^distal-K29^ and M1/K48/K63-linked triUb) and 80 μM WT Ub or Ub-K29R at 30°C for Ufd4-related ubiquitination or 37°C for TRIP12-related ubiquitination in the reaction buffer (50 mM HEPES, pH 7.5, 150 mM NaCl, 10 mM MgCl_2_ and 5 mM ATP). For the ubiquitination assays to validate key structural interfaces, the same concentrations of E1, E2, E3 enzyme, WT Ub, and fluorescent diUb^prox-K29^ were used.

To measure the ubiquitination activity on K48-linked substrate peptide and real N-end rule substrate, another ubiquitination assays were performed with 0.2 μM Uba1, 4 μM Ubc2, 0.25 μM Ubr1, 4 μM Ubc4, 0.25 μM Ufd4, 80 μM Ub, 5 μM (fluorescent degron, synthesized previously^36^) or 10 μM Scc1 (268–384) at 30°C with the same reaction buffer mentioned above.

These reactions were terminated by adding 4 × sodium dodecyl sulfate (SDS) sample buffer, and then analyzed using SDS-PAGE and visualized by Coomassie Brilliant blue dye/fluorescence or visualized by western blot using K29 linkage-specific antibodies^45^ and K48 linkage-specific antibodies (Abcam, ab190061).

### Kinetic measurement of Ub transfer on Ub and K48-linked diUb

To measure the ubiquitination velocity between Ub and K48-linked diUb, a time-resolved kinetic experiment was performed. *In vitro* ubiquitination assays were performed as described above, using fluorescent Ub-G76C or K48diUb as the substrate. The reaction mixtures were sampled and terminated by 4× SDS sample buffer at indicated time points (0, 1, 2, 5, 10, 15, 25, 45, 60 min), then analyzed using SDS-PAGE and imaged by fluorescence. Data from three replicates were quantified by image lab (Bio-Rad software) and time rate curves of ubiquitination were plotted.

### Transthiolation assay with fluorescently labeled Ub-MCQ

Transthiolation assay consists of two operational steps. First, 0.1 μM Uba1, 5 μM Ubc4 or its variants, 10 μM fluorescently labeled Ub-MCQ were mixed at 30°C for 5 min in the reaction buffer (50 mM HEPES, pH 7.5, 150 mM NaCl, 10 mM MgCl_2_ and 5 mM ATP) to generate fluorescent ubiquitin-loaded Ubc4 or its variants and then terminated with a final concentration of 250 mM EDTA (pH 7.4) on ice. Next, the loaded Ubc4 or its variants was reacted with Ufd4 or its variants to the following final concentrations: Ubc4 or its variants, 1.25 µM; fluorescent ubiquitin, 2.5 µM; Ufd4 or its variants, 0.5 µM and mixtures were sampled and terminated by 4× SDS sample buffer without Dithiothreitol (DTT) at 30 seconds, then analyzed using SDS-PAGE and imaged by fluorescence (Bio-Rad).

### Mass spectrometry analysis of polyubiquitin chains

To obtain polyubiquitin chains generated by Ufd4 or TRIP12 on K48-linked chains, *in vitro* ubiquitination assay was performed as described above and K48-linked tetraUb was used as a representative substrate. The reaction was quenched with 25 mM EDTA on ice. Then 5 μM engineered viral protease Lb^pro^* was added and incubated at 37°C overnight.

The identification and quantification of ubiquitin variants digested by Lb^pro^* were described as before^22^. In brief, diluted the reaction to 1 μM ubiquitin with a buffer (50 mM HEPES, pH 7.5, 150 mM NaCl). To perform high-mass protein analysis, samples were analyzed on a Q-TOF mass spectrometer (SYNAPT G2-Si, Waters company) instrument. The raw native electrospray mass spectra can be further deconvoluted by MaxEnt 1 (Waters) to generate a spectrum (relative intensity versus mass). This spectrum allows a single species to transform all the charge-state peaks into a single (zero-charge) peak. Microsoft Excel to further process the spectrum, and the intensities of un-(8,451.65 Da), single-(8,565.69 Da), and double-modified (8,679.73 Da) ubiquitin were exported.

To identify the K29 site and K48 site double-modified ubiquitin, the reaction digested after Lb^pro^* was further treated with a low concentration of wild type alpha-lytic protease (WaLP) at 4°C for 2 h. To analyze the specific linkage, a Thermo-Dionex Ultimate 3000 HPLC system was used and peptides were separated by a 60 min gradient elution at a flow rate of 0.3 μL/min. This system allows direct interface with the Thermo Orbitrap Fusion mass spectrometer. To identify peptide segments, the MS/MS spectra obtained from each LC-MS/MS run were analyzed using the Proteome Discoverer search algorithm (version 1.4), based on the target protein sequence. The search parameters were configured as follows: the mass tolerance for precursor ions was set to 20 ppm for all MS scans, and the mass tolerance for fragment ions was set to 0.02 Da for all acquired MS2 spectra. A fixed value PSM validator was used to estimate the peptide false discovery rate (FDR), and a peptide-spectrum match (PSM) was considered reliable if the q-value was below 1%.

### Isothermal titration calorimetry analysis

All ITC experiments were performed using a MicroCal PEAQ-ITC (MicroCal) in a buffer (50 Mm HEPES, 100 mM NaCl, pH 7.4) at 25 ℃. Purified Ufd4(164-706) fragment (28.5 μM) was titrated by K48-linked diUb (460 μM) or monoUb (407 μM), respectively. Data analysis and fitting were performed by MicroCal PEAQ-ITC Analysis Software.

### Cycloheximide Chase Analysis

For the Mgt1 degradation assays, ΔUfd4 yeast strain (a gift from Wei Li, Guangzhou Medical University) harboring the pRS313-Mgt1-6myc plasmid and the pRS316-Ufd4 plasmid or its variants were grown in SD (−His/Ura) medium containing 2% glucose (w/v) at 30℃ to an OD_600_ of 0.7 and then 68 μM MNNG was added to incubate for 40 min. Yeast cells were harvested by centrifugation and washed two times with sterile water. Cycloheximide at a final concentration of 1 mg/ml was added to the yeast cells and at each time point cells were collected and proteins were extracted as previously described^21^. Briefly, the pelleted cells were resuspended in 0.2 M NaOH for 5min at 25℃, then centrifuged to remove the supernatant and add 50ul loading buffer containing 8 M urea, 10% glycerol, 5% SDS, 1 mM EDTA, 100 mM DTT, 0.1% bromophenol blue and 0.2 M Tris-HCl, pH 6.8. The samples were heated at 70 °C for 8 min, followed by the supernatant after centrifugation was subjected to SDS-PAGE. The Mgt1 and GAPDH were detected by immunoblotting analysis using an anti-myc antibody (Cell Signaling Technology, 2272) and anti-GAPDH antibody (Proteintech, 60004-1-Ig), respectively.

### Generation of the probe capturing the transition state of the K29 linkage assembly on K48-linked diUb Preparation of Ub_(1–76)_-MesNa

The Ub_(1–76)_ hydrazine with a biotin tag at the N-terminus (1.0 equiv), prepared through protein chemical synthesis as previously described^23^, was dissolved in −15°C ligation buffer (2 mL, 100 mM NaH_2_PO_4_, 6 M Gn-HCl, pH 2.3). Then, NaNO_2_ (10 equiv) was added to the reaction, and the reaction was stirred for 30 min at −15°C. After that, 2-mercaptoethanesulfonate (MesNa, 50 equiv) was added to the reaction and the pH was slowly adjusted to 5.0-5.5. Next, the reaction reacted at room temperature for 3-4 hours. Finally, ligation buffer was added to dilute the reaction before purification, and the product was purified by semi-preparative HPLC using a Welch Ultimate XB-C4 column (21.2 × 150 mm, 5 μm particle size) and lyophilized.

### Preparation of K48 diUb-CAET-SAcm

1mM diUb^K29C^ with 40 mM (2-((2-chloroethyl)amino)ethane-1-(S-(Acetamidomethyl))thiol) were incubated in alkylation buffer (6 M Gn-HCl, 0.2 M HEPES, 10 mM TCEP, pH=8.0-8.5) at 30°C for overnight. The desired product was isolated by RP-HPLC and lyophilized.

### Preparation of K48 diUb-CAET-SH

Lyophilized K48 diUb-CAET-SAcm (final concentration 1 mM) was dissolved in reaction buffer (6 M Gn-HCl, 0.2 M NaH_2_PO_4_ buffer, pH 7.4). Next, fresh PdCl_2_ (15 equiv) was added to the reaction and the reaction was incubated at 30°C for 2-3 h. Finally, the reaction was quenched by adding DTT at a final concentration of 1 mM, and then the product was purified by semi-preparative HPLC and lyophilized.

### Preparation of K29/48 branched triUb-SH

Lyophilized K48 diUb-CAET-SH (1.0 equiv.) was dissolved in reaction buffer (6 M Gn-HCl, 100 mM NaH_2_PO_4_, Ph 7.4) to a final concentration of 1 mM. Next, 4-mercaptophenylacetic acid (MPAA, 50 equiv.) and Ub_(1–76)_-MesNa(1.5 equiv.) were added to the reaction buffer and the pH was adjusted to 6.5. Then, the reaction mixture was stirred at room temperature overnight, and the product was purified by semi-preparative HPLC and lyophilized.

### Preparation of K29/48 branched triUb probe

Lyophilized K29/48-triUb (1.0 equiv.) was dissolved in 1 mL buffer containing 6 M guanidinium chloride, 0.1 M NaH_2_PO_4_ buffer, pH 7.4 at a final concentration of 2 mg/mL. Then, 2,2′-Dipyridyl disulfide (AT_2_, 25 equiv.) was added to the buffer and the reaction mixture was stirred at room temperature for 2 h. Finally, the desired probe was purified and refolded with a Superdex75 column (GE Healthcare) pre-equilibrated with 50 mM HEPES, pH 8.0, 150 mM NaCl.

### Generation of the probe capturing the transition state of the Ufd4-Ubc4 transthiolation Preparation of Ub_(1–75)_-MesNa

The Ub_(1–75)_ hydrazine was prepared from Ub-G76C as previously described^46^ and was dissolved in - 15°C ligation buffer (2 mL, 100 mM NaH_2_PO_4_, 6 M Gn-HCl, pH 2.3). NaNO_2_(10 equiv) and 2-mercaptoethanesulfonate (MesNa, 50equiv) were then added to the reaction sequentially, as in the preparation of Ub_(1–76)_-MesNa. Finally, ligation buffer was added to dilute the reaction before purification, and semi-preparative HPLC and lyophilized purified the product.

### Preparation of Ubc4-CAET-SAcm

1mM Ubc4 mutant with only one Cys at C86 (Ubc4 C only) with 40 mM (2-((2-chloroethyl)amino)ethane-1-(S-(Acetamidomethyl))thiol) were incubated in alkylation buffer (6 M Gn-HCl, 0.2 M HEPES, 10 mM TCEP, pH = 8.0-8.5) at 30°C for overnight. The desired product was isolated by RP-HPLC and lyophilized.

### Preparation of Ubc4-CAET-SH

Lyophilized Ubc4-CAET-SAcm (final concentration 1 mM) was dissolved in reaction buffer (6 M Gn·HCl, 0.2 M NaH_2_PO_4_ buffer, pH 7.4). Next, fresh PdCl_2_ (15 equiv) was added to the reaction and the reaction was incubated at 37°C for 2-3h. Finally, the reaction was quenched by adding DTT at a final concentration of 1 mM, and then the product was purified by semi-preparative HPLC and lyophilized.

### Preparation of Ubc4-Ub-SH

Lyophilized Ubc4-CAET-SH (1.0 equiv.) was dissolved in reaction buffer (6 M Gn-HCl, 100 mM NaH_2_PO_4_, Ph 7.4) to a final concentration of 1 mM. Next, 4-mercaptophenylacetic acid (MPAA, 50 equiv.) and Ub_(1–75)_-MesNa(1.5 equiv.) were added to the reaction buffer and the pH was adjusted to 6.5. Then, the reaction mixture was stirred at room temperature overnight, and the product Ubc4-Ub was purified by semi-preparative HPLC and lyophilized.

### Preparation of Ubc4-Ub probe

Lyophilized Ubc4-Ub (1.0 equiv.) was dissolved in 1 mL buffer containing 6 M guanidinium chloride, 0.1 M NaH_2_PO_4_ buffer, pH 7.4 at a final concentration of 1.5 mg/mL. Then, 25 equiv. AT_2_ (2,2′-Dipyridyl disulfide) was added to the buffer and the reaction mixture was stirred at room temperature for 2 h. Finally, the desired product, Ubc4-Ub probe, was purified and refolded with Superdex75 column pre-equilibrated with 50 mM HEPES, pH 8.0, 150 mM NaCl.

### Sample preparation for single-particle cryo-EM

1.5 or 4.0 equiv K29/48 branched triUb probe, or Ubc4-Ub probe was mixed with Ufd4 (0.6 mg/mL) and incubated at 30°C in a water bath for 30 min. For TRIP12, 4.0 equiv K29/48 branched triUb probe was mixed with TRIP12 (0.6 mg/mL) and incubated at 25°C for 2 min. Prior to grid freezing using a Vitrobot mark IV (Thermo Fisher Scientific) running at 8°C and 100% humidity, 0.0001% fluorinated octyl maltoside (final concentration) was added to the samples. A volume of 3.5 µL sample was applied to glow-discharged Quantifoil Au 1.2/1.3 grids for 1 min and blotted for 1 s or 3 s, then immersed in liquid ethane to freeze.

### Cryo-EM Data collection and image processing

Cryo-EM micrographs were collected on a 300 kV Titan Krios microscope equipped with a Gatan K3 or Falcon 4 direct electron detector camera. Images were recorded using AutoEmation 2.0 at a pixel size of 1.074 or 0.808 Å with a total of 50 electrons over 32 frames. The defocus was set in a range of −1.5 μm to −2.0 μm. MotionCor2 (v. 1.2.6)^47^ and Gctf (v. 1.88)^48^ were used for motion correction and CTF parameter estimation. All the data were processed on RELION (v. 3.1)^49^ for particle picking, 2D classification, 3D classification, mask generation, refinement, postprocessing, and polishing. The detailed processing workflow can be found in Supplementary Fig. 21-23.

### Model building, refinement and validation

The cryo-EM maps of the Ufd4-triUb^K^^29/K48^ complex, TRIP12-triUb^K29/K48^ complex, and Ufd4/Ubc4-Ub complex sharped at different B-factors were used for model building. The AlphaFold predicted Ufd4 structure (code: AF-P33202-F1), AlphaFold predicted TRIP12 structure (code: AF-Q14669-F1-v4), ubiquitin (PDB code: 1UBQ) and Ubc4 (PDB code: 1QCQ) were fitted into the cryo-EM maps with rigid-body refinement in UCSF ChimeraX (v. 1.2.5)^50^. The individual models were merged into one molecule in WinCoot (V. 0.8.2)^51^. Real-space refinement in Phenxi (1.19.2)^52^ was subsequently carried out with secondary structure constraints and/or geometry restraints. Further iterative manual visualization and adjustment were conducted to give the final models. The statistics model building, refinement and validation were summarized in Extended Data Table S1. Structural analysis and figure layout were prepared in ChimeraX (v. 1.2.5).

## Data Availability

The cryo-EM maps and corresponding atomic coordinates generated in this study have been deposited in the Electron Microscopy Data Bank (EMDB) and Protein Data Bank (PDB) under accession codes EMD-35929 [https://www.ebi.ac.uk/emdb/EMD-35929], 8J1P (Ufd4-triUb^K29/48^) [https://www.rcsb.org/structure/8J1P]; EMD-35931 [https://www.ebi.ac.uk/emdb/ EMD-35931], 8J1R (Ufd4 in complex with Ubc4-Ub) [https://www.rcsb.org/structure/8J1R]; EMD-62292 [https://www.ebi.ac.uk/emdb/EMD-62292], 9KEN (TRIP12-triUb^K29/48^) [https://www.rcsb.org/structure/9KEN]. The MS/MS data generated in this study have been deposited to the ProteomeXchange Consortium via the PRIDE partner repository with the dataset identifier PXD062799 [https://www.ebi.ac.uk/pride/archive/projects/PXD062799]. Data for mass spectrometry characterization of semisynthetic proteins generated in this study are provided in Supplementary Fig.12-19. The atomic model of ubiquitin, Ubc4, Ub-loaded NEDD4 HECT domain, and E3-Ub-substrate complex are available under PDB accession codes 1UBQ [http://doi.org/10.2210/pdb1UBQ/pdb], 1QCQ [http://doi.org/10.2210/pdb1QCQ/pdb], 4BBN [http://doi.org/10.2210/pdb4BBN/pdb], and 4LCD [http://doi.org/10.2210/pdb4LCD/pdb]. The predicted atomic model of Ufd4 and TRIP12 are available in the AlphaFold Protein Structure Database under accession codes AF-P33202-F1 [https://alphafold.ebi.ac.uk/entry/P33202]and AF-Q14669-F1-v4 [https://alphafold.ebi.ac.uk/entry/ Q14669]. Newly created materials from this study may be requested from the corresponding authors. Source data are provided with this paper.

## Acknowledgments

We thank the National Natural Science Foundation of China (92253302, T2488301, 22137005, and 22227810 to L. L.; 22277073 and 92253302 to M. P.), National Key R&D Program of China (2022YFC3401500 to L. L.; 2023YFA0915300 to M. P.; 22407085 for Q.Z.), New Cornerstone Science Foundation (to L. L.). Shanghai Rising-Star Program (22QA1404900 to M. P.), Shanghai Pilot Program for Basic Research - Shanghai Jiao Tong University (21TQ1400224 to M. P.), Foundation of Muyuan Laboratory (118602240 to M. P.). Fundamental Research Funds for the Central University (to M. P.). Shanghai Jiao Tong University 2030 Initiative (WH510363003/003 to M. P., WH510363002/009 to Q. Z.), Shanghai Frontiers Science Center of Drug Target Identification and Delivery (ZXWH2170101 to H. A.), Chenguang Program of Shanghai Education Development Foundation (25Z520401224 to Q.Z.). We acknowledge the Tsinghua University Branch of China National Center for Protein Sciences (Beijing) for cryo-EM screening and data collection in 200 kV Arctica Tecnai microscopy and 300 kV Titan Krios microscopy. We also thank Shuimu BioSciences for Cryo-EM facility access and technical support during image acquisition.

## Author Contributions Statement

Conceptualization: X. W., H. A., J. M., Q. Z., L. Liu., and M. P. Design of experiments: X. W., H. A., J. M., Q. Z., L. Liu., and M. P. Synthesis of E2-to-E3 transthiolation intermediate mimics: J. M. Synthesis of E3-to-substrate ubiquitination intermediate mimics: X. W. and J. M. Cryo-EM sample preparation: J. M., X. W., and H. A. Structural determination of cryo-EM structures: H. A. and X.W. Structural modeling of the native transition states and chemically stable mimics: Z. D. Synthesis of fluorescently labelled K48-linked diUb and Ub: J. M., and X.W. Cloning and purification of E1, Ufd4, Ubc4, and all their mutants: J. M., X.W., H. C., L. Liang., and Z. T. Ufd4-related in vitro enzymatic assays and yeast assays: J. M., X.W., and H.C. Ufd4-related western-blot assays: Q. Z. and J. M. TRIP12-related in vitro enzymatic assays: X.W. Identification the topology of polyubiquitin chains generated by Ufd4 on K48-linked diUb: J. M. and Q. Z. The manuscript writing: X. W., H. A., J. M., Z. D., Q. Z., L. Liu., and M. P. The project supervision: Q. Z., L. Liu., and M. P.

